# Hidden sampling biases inflate performance in gene regulatory network inference

**DOI:** 10.64898/2025.12.19.695616

**Authors:** Marco Stock, Florin Ratajczak, Paul Bertin, Eva Hoermanseder, Yoshua Bengio, Jason Hartford, Pascal Falter-Braun, Matthias Heinig, Alexander Tong, Antonio Scialdone

## Abstract

Accurate reconstruction of gene regulatory networks (GRNs) from single-cell transcriptomic data remains a major methodological challenge. Recent machine learning approaches, particularly graph neural networks and graph autoencoders, have reported improved performance, yet these gains do not consistently translate to realistic biological settings. Here, we show that a key reason for that is the way negative regulatory interactions are sampled for supervised training and evaluation. We find that widely used sampling strategies introduce node-degree biases that allow models to exploit trivial graph-structural cues rather than biological signals. Across multiple benchmarks, simple degree-based heuristics match or exceed state-of-the-art graph neural network models under these biased evaluation protocols. We further introduce a degree-aware sampling approach that eliminates these artifacts and provides more reliable assessments of GRN inference methods. Our results call for standardized, bias-aware benchmarking practices to ensure meaningful progress in supervised GRN inference from single-cell RNA-seq data.

## 1 INTRODUCTION

Gene regulatory networks (GRNs) model the complex biological interactions that orchestrate gene expression. Revealing these regulatory interactions is key to understanding cellular function, dysfunction, and disease mechanisms. Therefore, accurate inference of regulatory networks from high-throughput genomic data is essential for unraveling the mechanisms that govern biological processes [1]. One natural approach to GRN inference is to leverage known regulatory relationships and train machine learning models in a supervised manner. In much of the recent literature, GRN inference is formulated as the reconstruction of the regulatory graph structure. In this setting, GRN inference is typically treated as a binary classification problem of inferring direct transcription factor (TF) binding events, using experimentally validated TF-DNA interactions, such as those derived from chromatin immunoprecipitation sequencing (ChIP-seq), as ground truth. Models are provided with a partially known regulatory graph together with single-cell RNA-seq (scRNA-seq) data, reflecting realistic biological scenarios in which known edges constitute only a small subset of existing interactions. Regulatory edges are commonly modeled as binary relationships, where the presence of an edge denotes a direct regulatory interaction [2, 3, 4].

For example, graph-based models frame GRN inference as a link-prediction task [5, 3, 2], where nodes represent genes and edges represent regulatory interactions between them. In particular, graph autoencoders (GAEs) learn low-dimensional embeddings of nodes for this purpose while integrating data on, e.g., gene expression levels as node features. By jointly encoding node features and known edges, GAEs can generalize to predict missing or uncertain regulatory interactions, making them well-suited for GRN inference. Beyond their theoretical appeal, several recent studies also report empirical performance gains in reconstructing GRNs [3, 2, 4].

GRN inference inherently operates under positive-unlabeled (PU) supervision: known regulatory interactions constitute the positive set, whereas the remaining gene pairs are unlabeled, not confirmed negatives. To construct a balanced test set of positive and negative examples of gene pairs for training and evaluation, GRN inference methods often rely on negative sampling, where a subset of unlabeled gene pairs is treated as negative examples, which is a special case of semi-supervised learning [6]. However, GRNs are extremely sparse, and the choice of negative sampling strategy strongly influences the statistical properties of the resulting dataset, thereby affecting model performance metrics such as AUROC (Area Under the Receiver Operating Characteristic curve) [7], which is widely used in recent benchmarking efforts [8, 4, 3, 2]. Topological benchmarking has assessed whether inference algorithms recover the structure of networks rather than only individual edges [9], yet how the choice of negative sampling impacts the performance evaluation has not been examined.

In this study, we show that negative sampling can introduce strong evaluation biases, which allows models to produce predictions based on simple graph-derived cues, such as node degree. It has already been shown on early ChIP-seq and gene expression datasets [10] that this causes evaluation pipelines to overestimate performance and obscure the true generalization ability of machine learning models. However, these findings have been largely ignored in single-cell GRN inference [11, 8] and in the resulting race in model development [8, 11, 3, 2]. We show that evaluation setups that ignore topological properties drastically inflate reported performance for link prediction in GRNs [3, 2]. The same sampling choices also affect methods that build GRNs from scRNA-seq data alone, as used in benchmarking frameworks such as BEELINE [8].

Our findings show that the negative sampling strategy is a central determinant of evaluation validity in GRN inference, and that carefully designed sampling is needed to avoid embedding spurious signal that models exploit for inflated accuracy. We therefore propose a degree-aware negative sampling strategy that removes degree-based structural bias. Applying this strategy to published models, we find that no method clearly outperforms random predictions, including advanced graph neural network approaches. Unbiased sampling is thus fundamental for a faithful assessment of both classical and machine learning based GRN inference methods.

Together, these results underscore the need for more rigorous and transparent evaluation frameworks in GRN inference. By identifying a key source of hidden bias and demonstrating its impact, this work establishes methodological principles that are broadly applicable across graph learning and systems biology research.

## 2 RESULTS

We first analyze commonly used negative sampling strategies in GRN benchmarking from a topological network perspective, and show how they distort key degree statistics. Based on this analysis, we propose an alternative, degree-aware sampling strategy that avoids these distortions. Second, we introduce the data and the models we evaluate, including variants that use the graph and expression data together or only one of the two, which lets us isolate their individual contributions. Third, we investigate how evaluation scores change across sampling protocols and expose strong biases. Finally, we show that non-learned predictors based only on node in- and out-degrees match or exceed published GAE models under their original evaluation settings.

### 2.1 Negative sampling strategies and their biases

Negative sampling plays a central role in GRN evaluation, because true negatives are unknown and models are trained and tested on artificially constructed negative edges. Which edges are plausible negatives is not a purely graph-theoretic question. Transcription factors (TFs) are proteins that bind DNA and regulate target genes, and they can be identified from their biochemical function and structural features, independent of any inferred network. Genes are therefore not interchangeable: only TFs can act as the source of a regulatory edge, whereas most genes can only be targets. A sampling scheme that ignores this, and, in effect, turns an arbitrary gene into a regulator by assigning it an outgoing edge, can produce biologically meaningless negatives.

We formalize this by representing the GRN as a directed, semi-bipartite graph *G* = {*V, E*} with directed edges *e*_*ij*_ = (*v*_*i*_ →*v*_*j*_), where the node roles follow from the ground-truth network together with this prior biology. A node acts as a TF if it has outgoing edges (out-degree *d*_*out*_(*v*_*i*_) *>* 0), as a target if it has incoming edges (*d*_*in*_(*v*_*i*_) *>* 0), and as isolated if it has neither; a node can be both TF and target. Even once node types are fixed a priori, a strong structural imbalance remains between high- and low-degree TFs, and it is this imbalance that a negative sampling strategy either respects or distorts. Published GRN inference tasks frequently use negative-sampling methods that we can broadly categorize into two types: random and structured sampling. Here, we also describe a third approach, degree-aware sampling, which selects negative edges while accounting for the degree distribution of nodes (Figure 1A).

**Figure 1.**
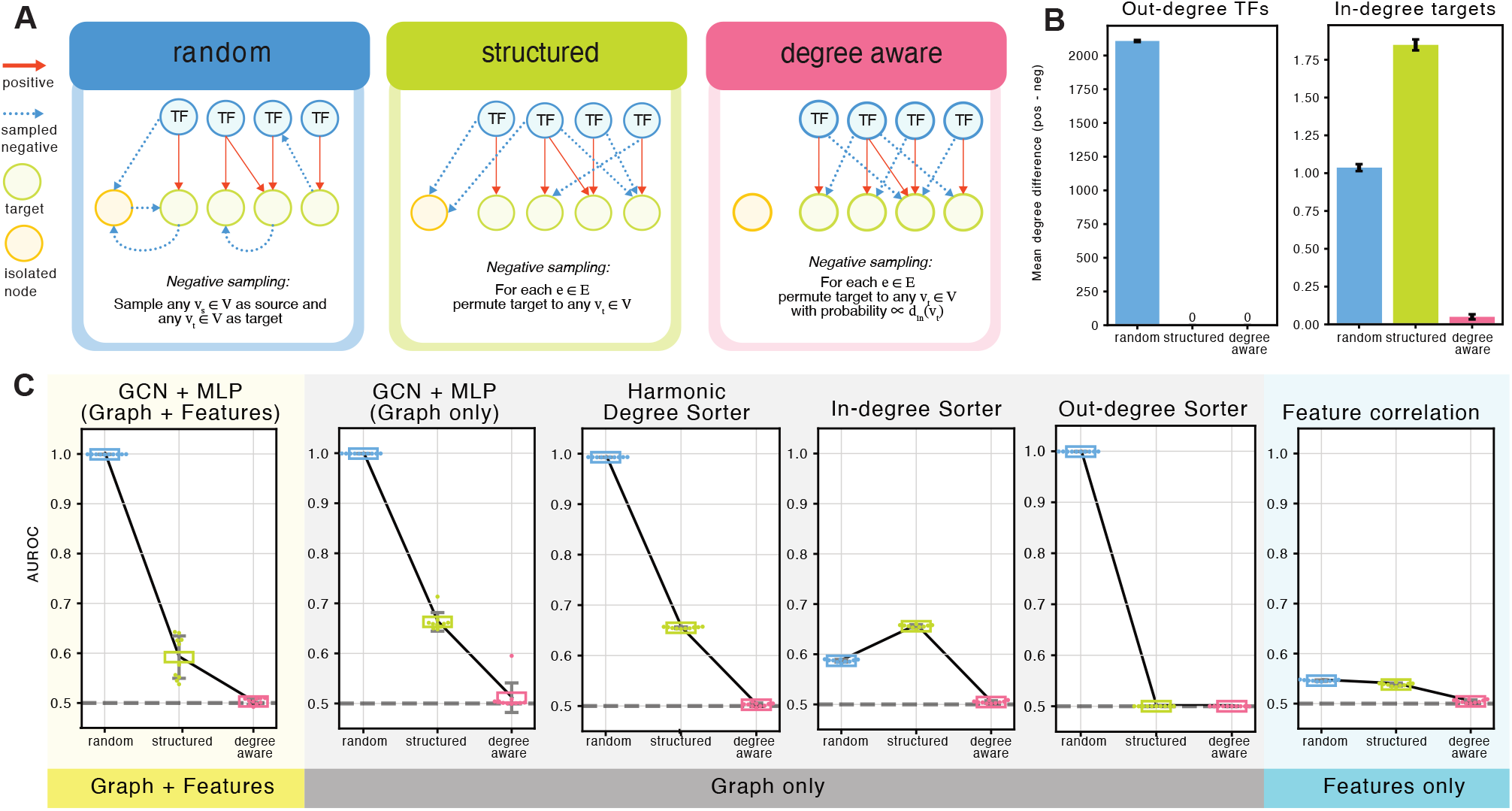
Effects of negative sampling on degree distributions and model performance. (A) Schematic representation and pseudo code for the negative sampling methods. (B) Mean difference between the out-degree of source nodes (left) and in-degree of target nodes (right) associated to positive and negative edges in the training graph. (C) Comparison of the Area Under the Receiver Operating Characteristic Curve (AUROC) values of various models with different negative sampling methods. We report these for models using training graph and scRNA-seq data (“Graph+Features”, left), only training graph (“Graph only”, middle), or only scRNA-seq data (“Features only”, right). TF = Transcription Factor; GCN = Graph Convolution Network; MLP = Multi-layer Perceptron. Results of additional “Graph+Features” models are available in Figure S1.

#### 2.1.1 Random Sampling

This approach randomly selects negatives uniformly over all possible edges without considering any underlying biological or structural information. Researchers often refer to this as global sampling, and they commonly employ it in large-scale benchmarking studies by evaluating the entire set of potential edges [8, 11], which effectively corresponds to random sampling with a sample size equal to the number of potential edges. Using the entire graph for every evaluation is computationally inefficient, as the number of potential edges scales quadratically with the number of nodes. Furthermore, this strategy introduces a severe imbalance between positive and negative edges, and this imbalance can bias performance metrics unless properly corrected [12]. In GRNs, this class imbalance translates directly into a structural bias: since most datasets contain relatively few TF nodes, random sampling predominantly selects negative edges that connect target genes to isolated or low-degree nodes. This produces pronounced differences in degree distributions, particularly in the out-degrees of TFs, between positive and negative edges (Figures 1B, S1A).

Importantly, GRNs are often hypothesized to follow a power-law degree distribution, in which a small number of TFs regulate many targets, while most TFs regulate only a few [13]. Random sampling disregards this structure, as it assigns equal probability to all edges, leading to a uniform expected node degree across sampled negatives. This mismatch between biological reality and sampling strategy can distort model evaluation and downstream interpretation.

#### 2.1.2 Structured Sampling

Structured sampling restricts the space of negative edges to a subset that is biologically or topologically closer to the positive edges. For each positive edge in the graph, we sampled a corresponding negative edge by corrupting one of its nodes. Several variants of this strategy exist, depending on which node is corrupted (source or target) and how replacements are selected. In our analysis, we adopted a commonly used variant in GRN inference [3, 2, 14, 15, 16, 4], where only the tail node is corrupted (i.e., the target gene). This ensured that all sampled negative edges originate from the same TF nodes as the positive edges. As a result, we preserved the out-degree distributions of TFs across both positive and negative edges (Figures 1B, S1B). This approach also avoids generating negative edges from isolated nodes, which would otherwise introduce specific topological artifacts (i.e., a peak at zero out-degree in the negative edge distribution that is absent from the positive edges; see Figure S1A) . By maintaining the TF-centric structure of the network, structured sampling provides a more biologically plausible and topologically consistent set of negatives for model training and evaluation. However, with this sampling procedure, there are still biases in the in-degree distribution, see Figure 1B.

#### 2.1.3 Degree-Aware Sampling

This strategy resembles structured sampling with the added feature that target nodes are sampled proportionally to their in-degree (Figure 1A). This additional constraint makes the marginal in- and out-degree distributions of negative edges comparable to those of positive edges, substantially reducing degree biases (Figures 1B, S1C). Because it matches the marginal degree distributions, it does not by construction equalize the joint (out-degree, in-degree) distribution or other topological features such as shared neighbors, path distance, or motif context, which we do not explicitly control.

Grover and Leskovec [17] previously introduced degree-based negative sampling in the node2vec method, where they bias node selection using a negative exponent to up-sample rare nodes. However, our approach differs in intent and implementation, as we aimed to preserve the natural degree distribution rather than counterbalance it.

Conceptually, degree-aware sampling resembles a subtype of configuration models, which are well-established in network theory and commonly used to study network motifs [18]. Despite its theoretical grounding, to our knowledge, no one has yet applied degree-aware negative sampling to the validation of recently developed scRNA-based GRN inference methods.

### 2.2 Methods and datasets used in GRN evaluation

While a continuously expanding selection of methods exists for GRN inference, many recent methods employ various flavors and combinations of graph autoencoders (GAEs) [2, 3, 15] and linear neural networks (i.e., multilayer perceptrons, MLPs) [16, 14, 15] for link-prediction tasks. Most of these models employ transductive learning [19], in which the objective is to predict the presence of edges in a partially observed GRN. To systematically assess how negative sampling influences the evaluation of such methods, we compared models that differ in the type of information they exploit: those that use both the graph prior and scRNA-seq features, those that use only the graph prior, and those that use only the gene expression features.

To compare methods on equal footing, we re-implemented the architectural components common to recent GRN inference methods within a single training framework and trained all models ourselves rather than reusing published weights. Each model has an encoder that maps its input (graph structure, expression features, or both) into a latent representation, and a decoder that predicts whether an edge *e*_*ij*_ exists from the embeddings *v*_*i*_ and *v*_*j*_ of genes *i* and *j*. Encoder and decoder are trained jointly as a binary classification problem. Using a single framework for all variants lets us attribute performance differences to only two factors, the information a model receives, and the negative sampling strategy used during training and evaluation. We evaluated models that use both the graph and expression data, that use only the graph, and that use only expression features.

First, we analyzed models that use both sources of data, i.e., the graph prior and the scRNA-seq features (“Graph + Features”). Expression levels for a given gene across all cells in a scRNAseq dataset form the initial node features fed to the model. The models include the GAE architectures commonly adopted in recent GRN-inference methods, as well as MLP-based link predictors. Together, they represent the set of graph-learning approaches that leverage expression-derived features while also exploiting topological cues from the partially observed GRN. These models are expected to achieve the best performance, since they can leverage both the graph prior and scRNA-seq data.

We then evaluated models that rely exclusively on the graph prior (“Graph Only”). Here, we include two distinct classes. First, we considered graph-learning models identical in architecture to the “Graph + Features” model but supplied an identity matrix as node features. This design eliminates expression-derived information and ensures that the model’s predictions depend solely on graph structure. Second, as negative controls that illustrate how negative sampling can induce simple but powerful topological biases, we include naive degree-based sorters that assign a score to each edge based only on the degrees of the incident nodes. These are not inference models, but they only exploit the topology and do not learn from biological patterns. Thus, we expect them to score close to a random predictor if the data contains no node-degree bias. The simplest variants estimate a score *ŷ* for an edge *e*_*ij*_ either by the out-degree of the source node *i* (out-degree sorter, ODS), or by the in-degree of the target node *j* (in-degree sorter, IDS):

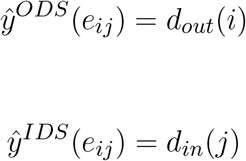

Because these methods are sensitive only to biases present in one of the degree distributions, we further included a combined approach: the harmonic mean sorter (HMS), which incorporates both endpoints of each edge:

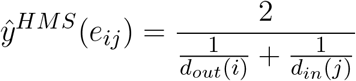

Finally, to provide a baseline that uses only scRNA-seq signal and no graph information, we included the simplest “Features Only” method: pairwise linear correlation that we computed directly from the expression matrix. This helps quantify how strongly the predictive signal from expression features alone is affected by the choice of negative sampling.

For our analysis, we used the largest scRNA-seq dataset curated by McCalla et al. [11] and originally published by Zhao et al. [20], which includes 36,199 mouse embryonic stem cells (mESCs) and 8,442 genes. ChIP-seq experiments provide the ground-truth GRN and the experimentally validated TF–target interactions (see Methods).

### 2.3 The sampling strategy affects evaluation

Here we show that, on the mESC dataset, model performance varies more with the negative-sampling protocol than with the model itself (Fig. 1C). This conclusion holds across multiple datasets and published methods (Table 1).

**Table 1.** AUROC values for published GCN-based methods (“Features + Graph”) and non-trained degree-based baselines (“Graph only”). Results are reported for the original training graphs provided by GNNLink [3] (“original”) and for graphs in which negative edges were resampled using a degree-aware strategy (“resampled”). Under the original evaluation setup, simple degree-based sorters (especially the “Out-Degree Sorter”) achieved performance comparable to or exceeding that of trained GCN-based methods. After degree-aware resampling, AUROC values for all methods collapse to near-random levels, indicating that the apparent performance gains are driven by degree-related sampling biases.

| Dataset | Features + Graph |  | Graph only |  |  |
| --- | --- | --- | --- | --- | --- |
|  | GNNLink [3] | GeneLink [2] | In-Degree Sorter | Out-Degree Sorter | H.-Mean Sorter |
| original |  |  |  |  |  |
| hESC 500 | 0.823 | 0.810 | 0.647 | **0.844** | 0.751 |
| hESC 1000 | 0.837 | 0.831 | 0.633 | **0.853** | 0.731 |
| mESC 500 | 0.833 | 0.851 | 0.661 | **0.864** | 0.841 |
| mESC 1000 | 0.821 | **0.897** | 0.671 | 0.863 | 0.836 |
| resampled |  |  |  |  |  |
| hESC 500 | **0.517** | 0.497 | 0.500 | 0.500 | 0.500 |
| hESC 1000 | **0.523** | 0.504 | 0.500 | 0.500 | 0.500 |
| mESC 500 | **0.522** | 0.498 | 0.500 | 0.500 | 0.500 |
| mESC 1000 | **0.519** | 0.501 | 0.500 | 0.500 | 0.500 |

Under random sampling, all models with access to the graph prior, including the GCN-based as well as the degree-based sorters, achieve near-perfect AUROC values (see Figure 1C and S1D). This inflated performance is expected, as random sampling generates negative edges predominantly involving low-degree or isolated nodes, producing a strong degree contrast between positive and negative examples (Figure 1B). Therefore, because of this bias, any model that can read out degree performs extremely well, regardless of whether it captures meaningful biological relationships.

Under structured sampling, performance decreases for all graph-informed methods but remains artificially elevated relative to a setting without degree bias. This is because structured sampling preserves the out-degree distribution of TFs across positive and negative edges but leaves the in-degree distribution of target genes unbalanced (Figure 1B and S1B). As a result, even simple models like in-degree-based heuristics continue to exploit residual structural cues. Consistent with this, the out-degree sorter drops to random-level performance, while the in-degree sorter and graph-based models retain substantial predictive power (Figure 1C and S1E). As shown in Figure S1E, the comparison between models under structured sampling reveals slight performance benefits of the trainable MLP decoder compared to the fixed decoding that uses the inner product of the embeddings, which can be expected from the larger capability and complexity of the trainable model.

Only under degree-aware sampling the degree-based shortcuts disappear. By matching both in-degree and out-degree distributions between positive and negative edges, degree-aware sampling eliminates the topological asymmetries that random and structured sampling introduce (Figure 1B-C and S1C). Consequently, all models that rely exclusively on graph structure, including degree sorters and graph-only GCN variants, fall to near-random performance (Figure 1C and S1F).

Once these within-method patterns are clear, we can meaningfully compare the methods to each other. Under both structured and degree-aware sampling, GCN models that use only graph structure perform as well as (or sometimes better than) their counterparts that also incorporate scRNA-seq features. This suggests that the feature-derived signal is weaker than the strong structural bias present under commonly used sampling strategies. Feature-only correlation baselines also drop substantially under degree-aware sampling, indicating that feature similarity itself correlates with graph connectivity in the training graph and is therefore indirectly influenced by the same biases.

One might argue that subsampling negatives is itself the problem, and that methods should be scored on the complete adjacency matrix. However, evaluating on the unsampled graph carries a bias comparable to random sampling (Figure S3). When predictions are scored on all candidate pairs, the global (micro) AUPRC reflects the same marginal degree distribution that random sampling reproduces, and it rewards the same out-degree shortcut exploited by the graph-only GCN (Figure S3A). Computing AUPRC within each TF and averaging across TFs (a per-regulator evaluation of the kind advocated by McCalla et al. [11]) holds the source node fixed and removes the out-degree shortcut, so the GCN and the out-degree sorter both fall to the random baseline. The in-degree bias remains, and the in-degree sorter stays well above random (Figure S3B). This is the same residual we observed under structured sampling, which balances the out-degree but not the in-degree distribution. Scoring on the full graph therefore does not deliver the unbiased comparison that structured sampling provides.

Overall, these results show that (i) negative sampling choices dominate evaluation outcomes, (ii) performance differences between models largely vanish once degree bias is removed, and (iii) the apparent superiority of recent GAE-based approaches might arise primarily from biased sampling rather than from genuine improvements in GRN reconstruction.

### 2.4 Sampling biases underlie reported gains in recent GRN inference models

The sampling-induced biases described above are not limited to controlled benchmarking scenarios, but they also affect results reported in the literature. To assess the extent of this issue, we revisited widely used evaluation datasets released with GeneLink [2] and reused by other GRN-inference methods, including GNNLink [3] and GT-GRN [4]. These sets are based on data first published in BEELINE [8] and provide both the positive edges and the pre-sampled negative edges used to compute the published performance values, allowing us to directly examine whether structural biases are present in their evaluation protocol. We found that the negative edges included in these datasets exhibit pronounced imbalances in both in-degree and out-degree distributions (Figure S2), consistent with the biases introduced by the sampling strategies analyzed above. As a result, simple degree-based heuristics, which exploit these imbalances but contain no biological or learned information, achieve performance that matches or even exceeds that of original machine learning models (Table 1). For example, the out-degree sorter consistently outperforms GNNLink and GeneLink across all datasets. Replacing the originally published negative edges with edges sampled using the degree-aware sampling regime drastically reduces the performance of both the naive and published models (Table 1), demonstrating that the high AUROC values originally reported can be explained by topological shortcuts rather than genuine regulatory learning. This analysis confirms our central conclusion, which rests on the resampled comparison. Once the negative edges are redrawn with degree-aware sampling, the degree contrast that the naive predictors exploit is removed, and both naive and trained methods fall to near-random performance (Table 1).

These findings show that sampling bias is not merely a theoretical concern: it directly affects published claims of improved GRN-inference accuracy. Because the datasets supplied by previous studies encode their own degree-based artifacts, any method evaluated on them, regardless of its architectural sophistication, is granted access to shortcuts that may artificially inflate measured performance. This highlights the need to incorporate bias-aware negative-sampling procedures as a necessary component of GRN benchmarking, to ensure that reported improvements reflect true predictive ability rather than dataset-specific structural artifacts.

## 3 DISCUSSION

The results presented here demonstrate that negative sampling strategies play a central and often underappreciated role in the evaluation of GRN inference methods. By systematically comparing random, structured, and degree-aware sampling, we show that AUROC values commonly reported in recent GRN studies are dominated by sampling-induced degree imbalances rather than by a model’s ability to learn regulatory relationships. When degree biases are removed, graph-based models, graph-only variants, and naive degree heuristics all converge to near-random performance. The differences between these methods under standard benchmarking therefore reflect properties of the sampling protocol and the ability of models to exploit them, rather than genuine differences in modeling capability. In particular, GAE-based models experience a sharp decline in model performance from almost perfect to almost random, both when comparing standard models on data from McCalla *et al*. [11] (Figure 1C), as well as comparing published methods [2, 3] on datasets that were originally published in [8] and have been reused across recent GRN inference method development [2, 3, 4] (Table 1). Importantly, even the method based only on feature correlation, which has no direct access to degree information, drops in performance once degree-aware sampling is used (Figure 1C), indicating that the chosen evaluation setup can also influence benchmarking results of feature-only methods, such as those presented in BEELINE [8] and Mccalla *et al*. [11], effectively questioning the reported performance of many methods developed over the last five years.

This decline in correlation-based approaches suggests that conventional benchmarks may obscure signal limitations that are independent of model architecture and that degree-aware sampling serves as a diagnostic tool to investigate what models have actually learned, regardless of their complexity. This does not imply that evaluations based on conventional sampling are without merit. Our point is rather that such protocols leave degree structure in the negative set, and that models are quite capable of learning it; apparent gains may therefore index that capacity rather than real progress in inference. Degree-aware sampling provides a complementary, stricter setting and a necessary standard against which claims of progress in GRN inference should be assessed.

Why do such degree-based shortcuts arise so easily? GRNs exhibit strongly heterogeneous degree distributions: a small number of TFs regulate many genes, whereas most regulate only a few. This pattern is reminiscent of preferential-attachment processes, in which nodes with more edges attract additional connections [21, 22, 23]. Although the exact generative mechanisms underlying GRN topology remain debated [24, 23], high-degree regulators are well documented in empirical TF-target maps. When negative edges are sampled without accounting for this heterogeneity, the resulting imbalance allows models to perform well simply by predicting edges involving high-degree nodes. Our findings, therefore, do not rely on assuming a strict power-law distribution, but they do highlight how features often associated with preferential-attachment-like structures create vulnerabilities in training and evaluation if not properly controlled.

What we observed is a domain-specific instance of shortcut learning, in which models exploit superficial cues that are predictive on a benchmark but do not reflect the intended task [25]. GNNs are especially susceptible, as prior work has shown that they tend to over-exploit degree information [26]. The most advanced graph-based models are, thus, also the ones most prone to leveraging sampling-induced shortcuts instead of learning biologically meaningful regulatory patterns. Our results show this also for methods based on Graph Attention Networks (GAT), which are not immune to the issue (Figure S1): attention mechanisms can refine how information is aggregated across the graph, but they cannot correct for shortcuts encoded in the training data itself. Architectural sophistication is therefore no substitute for unbiased evaluation.

Beyond GRN inference, negative sampling design has received substantial attention in the broader graph representation learning community, where increasingly sophisticated strategies have been proposed [27, 28]. These include adversarial approaches such as KBGAN [29] and Markov chain-based samplers, which aim to generate negative edges that closely resemble true positives to reduce sampling-induced artifacts and improve downstream task performance [27, 28]. Graph representation learning research has also repeatedly shown that GNNs tend to overemphasize high-degree nodes, a bias documented across dozens of studies [30], with normalization schemes proposed to mitigate degree-related effects [31, 32]. While these advances offer promising directions, their effectiveness for GRN inference remains untested, and our results indicate that even simple degree-aware sampling is sufficient to eliminate the most severe biases caused by degree imbalance.

Our findings should not be taken to imply that degree heterogeneity is biologically meaningless. High-degree regulatory hubs are a real feature of GRNs. The point is specific to evaluation. Node degree is computed from the observed graph and is therefore information a model is given, not a quantity it must infer. An evaluation in which positive and negative edges are separable by degree consequently rewards reproducing a known input rather than identifying which specific TF-target pairs interact. In the limiting case where degree alone determined the labels, no inference algorithm would be needed at all. Removing the degree contrast between positives and negatives therefore discards no biological signal that a GRN method should be credited for but it withholds information the method already possesses. In this context, recent GRN inference approaches based on generative formulations are also relevant, because they can reduce or avoid explicit negative sampling and thus may help clarify whether the performance decline observed here is specific to supervised edge-classification pipelines or reflects a broader limitation of the available input signal.

Another approach to improve negative sampling for GRNs requires additional knowledge in the form of disease-associated genes [33], which is itself an ongoing field of research with its own problematic biases [34]. Patterns similar to those we report have been observed in other biological graph-learning settings, where uniform negative sampling inflates performance on biomedical knowledge graphs [35, 36, 37] and protein interaction prediction [38], and degree-related artifacts have been shown to drive misleading accuracy estimates across various domains [39, 40]. In this work, we show that such biases are present in datasets that form the foundation of recent GRN inference method development and we quantify the extent to which model performance is driven by biased degree distributions. This is especially crucial since recently developed GRN inference methods continue to employ negative sampling strategies that expose models to these shortcuts [4].

These observations have several implications for future GRN inference method development. First, benchmarking frameworks should incorporate negative-sampling procedures that explicitly balance both in- and out-degree distributions, such as the degree-aware strategy introduced here. This is the minimal step required not only to prevent models from exploiting trivial topological cues, but also to quantify the extent to which existing methods rely on them. Second, evaluations should include graph-only baselines to assess whether performance gains arise from biological signal or from graph structure alone. Third, since the performance decline under degree-aware sampling may partly reflect a broader limitation of expression-derived signal not specific to supervised graph models, evaluation frameworks should include settings that do not rely on explicit negative sampling, such as generative model formulations. This would help assess whether performance gains reflect genuine improvements in regulatory signal capture, independent of how negatives are constructed. External validation using independently generated datasets [10] or perturbation-based assays remains essential for determining whether inferred networks generalize beyond dataset-specific artifacts.

Our analysis establishes this mechanistic picture in depth on a large mESC dataset, and the re-evaluation of published methods across four additional datasets shows that the same degree artifacts appear in widely used benchmarks. A natural extension is to replicate the full sampling-by-model comparison across more datasets and network types, which would map how the balance between sampling and model choice shifts with network density, TF/target ratio and species. Because training and evaluation negatives are matched in each run, our design cannot separate the two. Crossing these conditions would establish whether the effect arises primarily during training or during evaluation.

In summary, our results reveal a fundamental methodological challenge in supervised GRN inference: sampling-induced biases, amplified by natural degree heterogeneity in regulatory networks, can produce misleading performance estimates for complex models. By adopting bias-aware sampling strategies and more stringent benchmarking practices, the field can obtain more reliable estimates of method performance and accelerate genuine progress in reconstructing GRNs from single-cell data.

## 4 METHODS

### 4.1 Datasets

The dataset used was obtained from McCalla et al. [11]. We chose the largest available dataset from their study, which is a mouse embryonic stem cell dataset (mESC) with 36,199 cells and 8,442 genes from [20]. As our ground truth, the provided ChIP-Seq derived GRN was used.

After removing self-loops, the ground truth network was split into a non-overlapping training, validation and test set of interactions by randomly sampling positive edges in a ratio of 60/20/20 percent. The negative edges were then sampled for training, validation and test set by using the respective negative sampling technique. The negative training edges were checked to not overlap with the positive training edges, but could include positives of validation and test set, since removing these from the training set would be considered an information leak to the training set. The validation negative edges were checked to not overlap with validation or training positives. The test negative edges were checked to not overlap with any training, validation and test positives. Since this approach is a semi-supervised approach in a positive-unlabeled setting, the negative sampling was always done for a set of positive edges and did not take previously sampled negatives into account.

The preprocessed scRNA-Seq data provided was min-max scaled for each gene across cells.

### 4.2 Reevaluation of GeneLink dataset

Pre-sampled positive and negative edges used to calculate the AUROC scores shown in Table 1 are obtained from [3]; they are based on cell-type-specific ChIP-seq datasets provided in the BEELINE benchmarking study [8], and have also been used in [2] and [4]. Training, validation and test splits were downloaded from https://github.com/sdesignates/GNNLink/tree/master/Data/Train_validation_test. Performance of naive models is reported on the test set, using the degrees of the training and validation sets jointly. Performance on trainable models GNNLink [3] and GENElink [2] was recalculated using their original published code and the respective model’s default parameters. For the resampled performance, we kept the positive edges in training, validation and test set identical to the original and only replaced the negative edges using the degree-aware sampling strategy. Since both models had the tendency to overtrain on the resampled splits, we reduced the number of training epochs to 20 to allow a more realistic comparison of performance. Performance of [4] was not included in the comparison since retraining the model requires additional data that is not published.

### 4.3 Negative Samplings

For random sampling the *negative_sampling* function of the torch_geometric python package [41] was used. For structured sampling the *structured_negative_sampling* function of the same package was used. The degree-aware sampling was implemented manually, see Data and Code Availability Statement.

### 4.4 Models

#### 4.4.1 Graph + Feature models

We benchmarked the performance of different combinations of graph autoencoder models. Each graph autoencoder consists of an encoder, which takes the input data and projects it into a latent space, and the decoder, which takes the latent embeddings *v*_*i*_ and *v*_*j*_ of genes *i* and *j* and predicts if the edge *e*_*ij*_ is present in the data as a binary classification task. Trainable parameters of encoder and decoder were optimized jointly. The reconstruction loss was minimized during training, as measured by the binary cross entropy loss across training edges. For the encoder, we evaluated graph convolutional layers (GCN), as well as graph attention layers (GAT). For the decoder, we evaluated the standard non-trainable GAE decoder, ranking interactions based on the InnerProduct between the embeddings of each gene pair, as well as a trainable MLP decoder.

#### 4.4.2 Graph-only baseline and degree-based negative controls

To assess the performance achieved by models that rely only on graph training data, two different types of models were used. Naive models are static, not trainable functions, ranking edges based on: the in-degree of the target node (“In-degree sorter”), the out-degree of the source node of an edge (“Out-degree sorter”) and a third model ranking edges based on the harmonic mean of these two values (“Harmonic degree sorter”). We complemented the non-trainable naive models by a graph autoencoder that has only the training edges as inputs. The scRNA-Seq feature matrix was replaced with the identity matrix, leading to one-hot encoded identity vectors for the nodes as node features. This model uses GCN layers for message passing in the encoder and an MLP for the decoder.

#### 4.4.3 Feature only baseline

As a baseline for the performance achieved from only features from scRNA-Seq data, the linear correlation between the gene expression profiles of each pair of genes along the different cells was used to score the edges.

#### 4.4.4 Model evaluation

For each trainable model we performed a hyperparameter optimization using bayesian optimization to maximize the area under the precision-recall curve on the validation set. This was done separately for each sampling type. For the optimization, we considered the following parameters:

| Parameter | Distribution | Values / Range |
| --- | --- | --- |
| dropout | uniform | [0, 1] |
| layer ratio (encoder) | integer uniform | [5, 20] |
| learning rate | uniform | [0.0001, 0.01] |
| epochs | discrete | {200, 500, 1000} |
| early stopping patience | discrete | {5, 10, 20} |
| latent dimensions | discrete | {16, 32, 64, 128, 256} |
| number of convolution layers | discrete | {1, 2, 3} |

We then chose hyperparameters that achieved the best validation AUROC performance on the respective validation set of up to 250 runs.

Afterward, we report ten different test scores of ten models trained with the selected hyperparameters for different seeds for the split of training, validation and test edges. The type of negative sampling used for training and evaluation is always matching. For the non-trainable models, we directly report the results of these models on the same 10-fold set of different seeds.

## AUTHOR CONTRIBUTIONS

M.S. and F.R. generated, analyzed data, produced the figures. M.S., F.R., P.B., J.H., A.T. and A.S. interpreted the results. M.S. and A.S. conceived the study. P.F.B., M.H., E.H., Y.B., J.H., A.T. and A.S. provided project supervision. M.S., F.R, and A.S. wrote the manuscript.

## ACKNOWLEDGMENTS

M.S. and F.R. were supported by the Helmholtz Association under the joint research school “Munich School for Data Science - MUDS”. M.S. was supported by a Joachim Herz Stiftung Add-on Fellowship for Interdisciplinary Life Science. Work in the Scialdone lab is supported by the Helmholtz Association and the DFG (Project number 448727785). The lab of Y.B. acknowledges funding from UNIQUE, CIFAR, NSERC, Intel, and Samsung. M.H. is supported by the Chan Zuckerberg Foundation (2019-202666, 2021-237882) and the DZHK (German Center for Cardiovascular Research) projects 81Z0600106 and 81Z0600105. Work in the P.F.B. lab was supported by the Free State of Bavaria’s AI for Therapy (AI4T) Initiative through the Institute of AI for Drug Discovery (AID), and the Deutsche Forschungsgemeinschaft (DFG) Project FA 1462/5-1 .

## DATA AND CODE AVAILABILITY STATEMENT

All code used in this study, including the code to generate the figures, is available on GitHub (https://github.com/ScialdoneLab/Reevaluating-GRN-inference). All data logged during model training and hyperparameter optimization are available on Weights and Biases (https://wandb.ai/scialdonelab/GRN_inference). The positive and negative edges used to compute the metrics reported in Table 1 were obtained from https://github.com/sdesignates/GNNLink.

## SUPPLEMENTAL DATA

**Figure S1.**
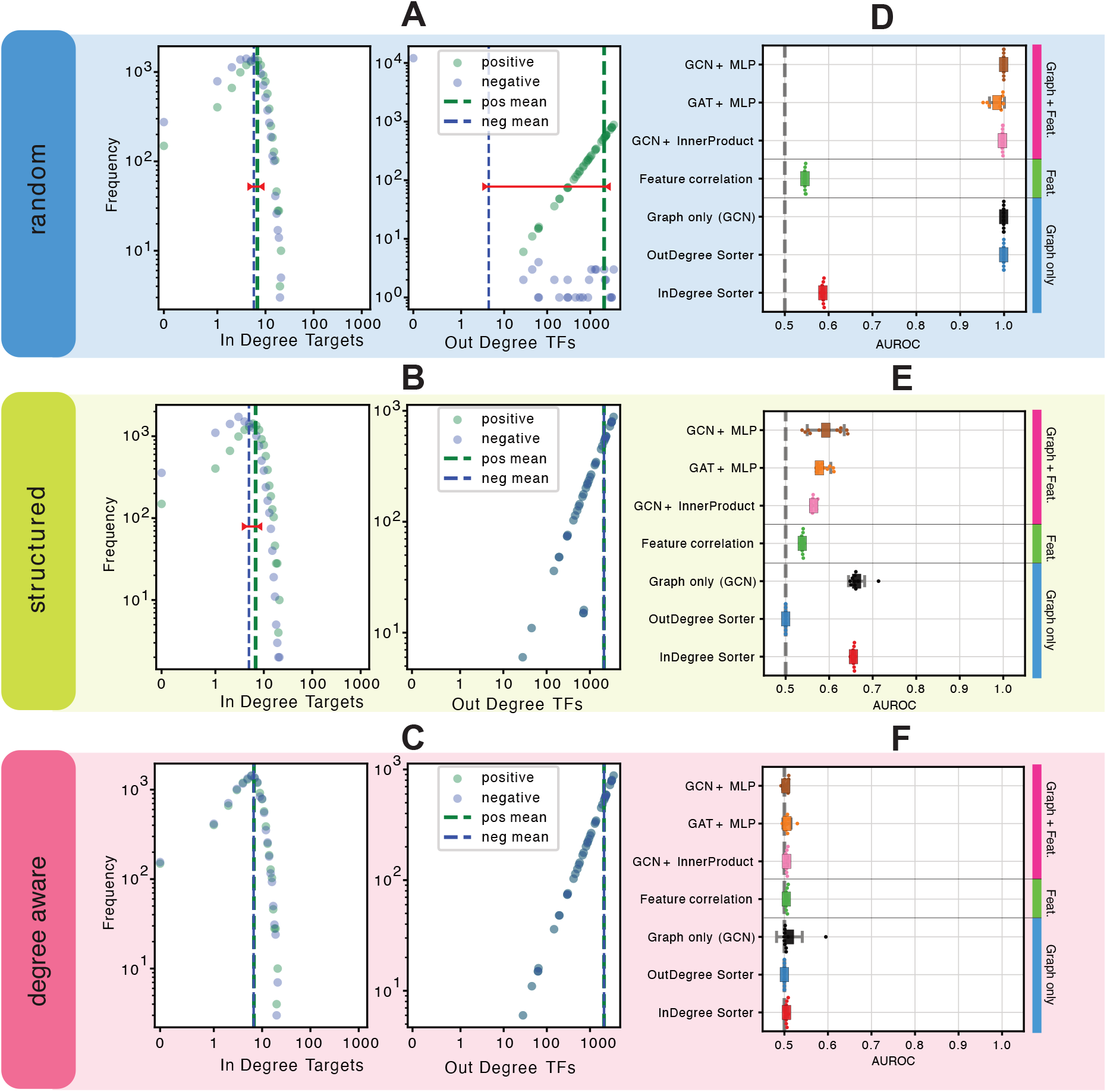
Comparison of different sampling methods for negative edges in the ground truth graph (A)-(C) Node degree distribution of in-degrees target nodes (genes) and out-degrees of source nodes (transcription factors) for edges in test set with different negative sampling methods. Degrees plotted are derived from the encoding graph used for test prediction, which includes all positive training and validation edges. Vertical line denotes mean degree of the respective degree distribution. Horizontal red line denotes difference in mean degree between positive and negative edges. (D)-(F) Comparison of the evaluation results of different models using the different negative samplings.

**Figure S2.**
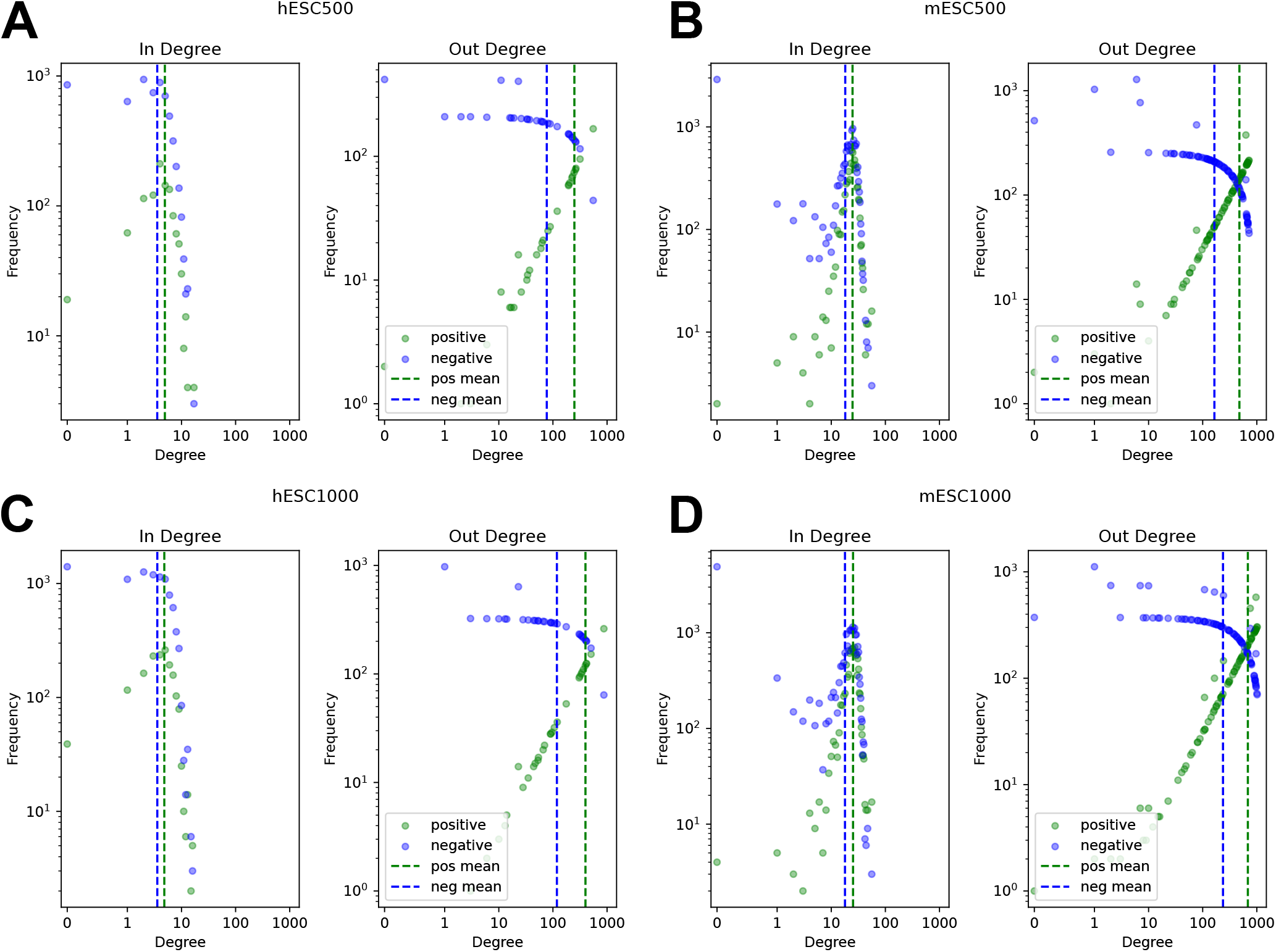
Node degree distributions of published positive and negative edge sets for four datasets used in [2, 3, 4]. Shown are the node degrees of published test edges based on a graph that is build from combining the published training and validation edges.

**Figure S3.**
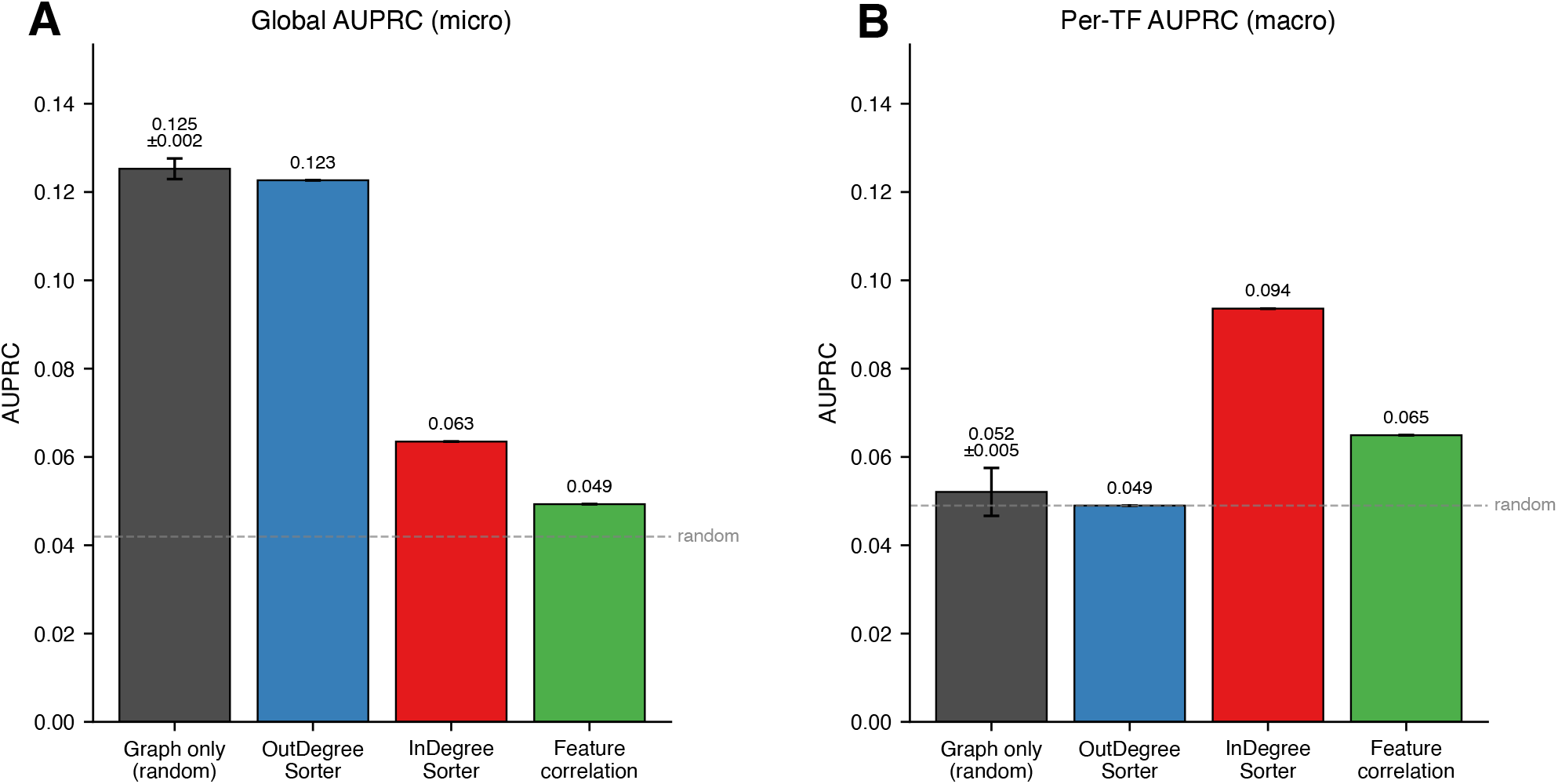
AUPRC for predictions on the unsampled graph of a graph-only GCN+MLP autoencoder trained under random negative sampling, the out-degree and in-degree sorters, and the feature-correlation baseline. (A) Global AUPRC on the whole adjacency matrix. (B) Per-TF (macro) AUPRC, computed within each TF’s candidate targets and averaged over TFs. Dashed lines show random expectation. Autoencoder bars show mean ± standard deviation over 10 cross-validation folds with different train/val splits, but fixed test set. The degree sorters and feature correlation are deterministic on the fixed test set.

